# Multiple mTOR RNA localization signals regulate subcellular protein synthesis and axonal growth

**DOI:** 10.64898/2026.03.22.713398

**Authors:** Nitzan Samra, Pabitra K. Sahoo, Agostina Di-Pizio, Courtney N. Buchanan, Nataliya Okladnikov, Ofri Abraham, Shifra Ben-Dor, Rebecca Haffner-Krausz, Ida Rishal, Jeffery L. Twiss, Mike Fainzilber

## Abstract

Subcellular localization of mTOR is thought to be key for regulating cell size and growth, but the relative contributions of mRNA versus protein localization are unclear. We used reporter mRNA localization assays to identify two distinct mTOR Localizing Sequences (MLS) in its 5’UTR, in addition to the localization activity already reported for the 3’UTR. Gene-edited mice with deletion of both 5’UTR MLS are mTOR hypomorphs with reduced body weight and brain size. In contrast, a mouse line lacking the second 5’UTR MLS and the 3’UTR retains near normal overall mTOR expression levels with specific subcellular perturbation of mTOR localization to neuronal axons. This subcellular mTOR deficit affects axonal local protein synthesis and neuronal growth. Thus, mTOR transcripts are localized by multiple UTR sequences, and subcellular localization of mTOR mRNA regulates local protein synthesis and neuronal growth.

## Introduction

mTOR is a master regulator of cellular metabolism, protein synthesis and cell growth [1, 2]. Subcellular localization and transport of mTOR protein are thought to be critical for its functions [3–5]. Distinct mTOR complexes associated with lysosomes or endomembranes in diverse cdell types and axonal mTOR in neurons respond to local cues and regulate adjacent translation events [4–6]. RNA localization provides an efficient means for subcellular regulation of signaling events [7, 8], but the relative contributions of protein versus RNA localization to mTOR functions are not well understood.

mTOR promotes growth of both central and peripheral neurons [9, 10], and the extended morphology of mature neurons provides an advantageous model for the study of localization mechanisms. Our previous work has identified intrinsic neuronal growth regulation by local translation of kinesin transported mRNAs in the axon and retrograde transport of the newly synthesized proteins in a complex centered on importins and dynein [11–14]. We and others have shown that mTOR mRNA is localized to axons [15–19] and the mTOR 3’UTR was shown to have axon localization activity [16]. However, mTOR 3’UTR deletion neurons retained a degree of mTOR expression in axons and did not show significant growth effects [16].

We asked if additional axonal localization determinants are present in mTOR mRNA and if so, what are their functional effects? Two distinct axonal localization elements were identified in the short mTOR 5’UTR, in addition to the localization activity previously reported in the 3’UTR. Gene-edited mice with deletion of both 5’UTR mTOR axonal localization elements presented as mTOR hypomorphs, while mice with deletions of the second 5’UTR element together with the 3’UTR presented axonal mislocalization of mTOR mRNA without effects on somatic mTOR levels. The latter mutant revealed subcellular protein synthesis and neuron growth phenotypes.

## Results

### Two mRNA localization motifs in the mTOR 5’UTR

As noted above, the mTOR 3’UTR deletion mouse retained approximately 50% of wild type levels of axonal mTOR mRNA [16]. Since axonal localization of mRNAs can also be mediated by 5’UTR sequences [20–23], we used single molecule fluorescence in situ hybridization (smFISH) to test axonal localization capacity of the mTOR 5’UTR fused to an EGFP reporter mRNA. The full 5’UTR conferred effective axonal localization of the reporter mRNA (Figure 1A, B). Testing of partial segments of the UTR as indicated in Figure 1A led to the identification of two distinct 5’UTR mTOR localization signals (MLS) (Figure 1B, C and Extended Figure 1A). Both MLS sequences are highly conserved across vertebrates (Figure 1D-F), with sequence identities of 87.2% for MLS1 and 95.6% for MLS2 between mouse and human. There is little homology between the two MLS sequences, however both are predicted to form stem-loop structures containing guanine-rich sequence stretches (G-stretches).

**Figure 1.**
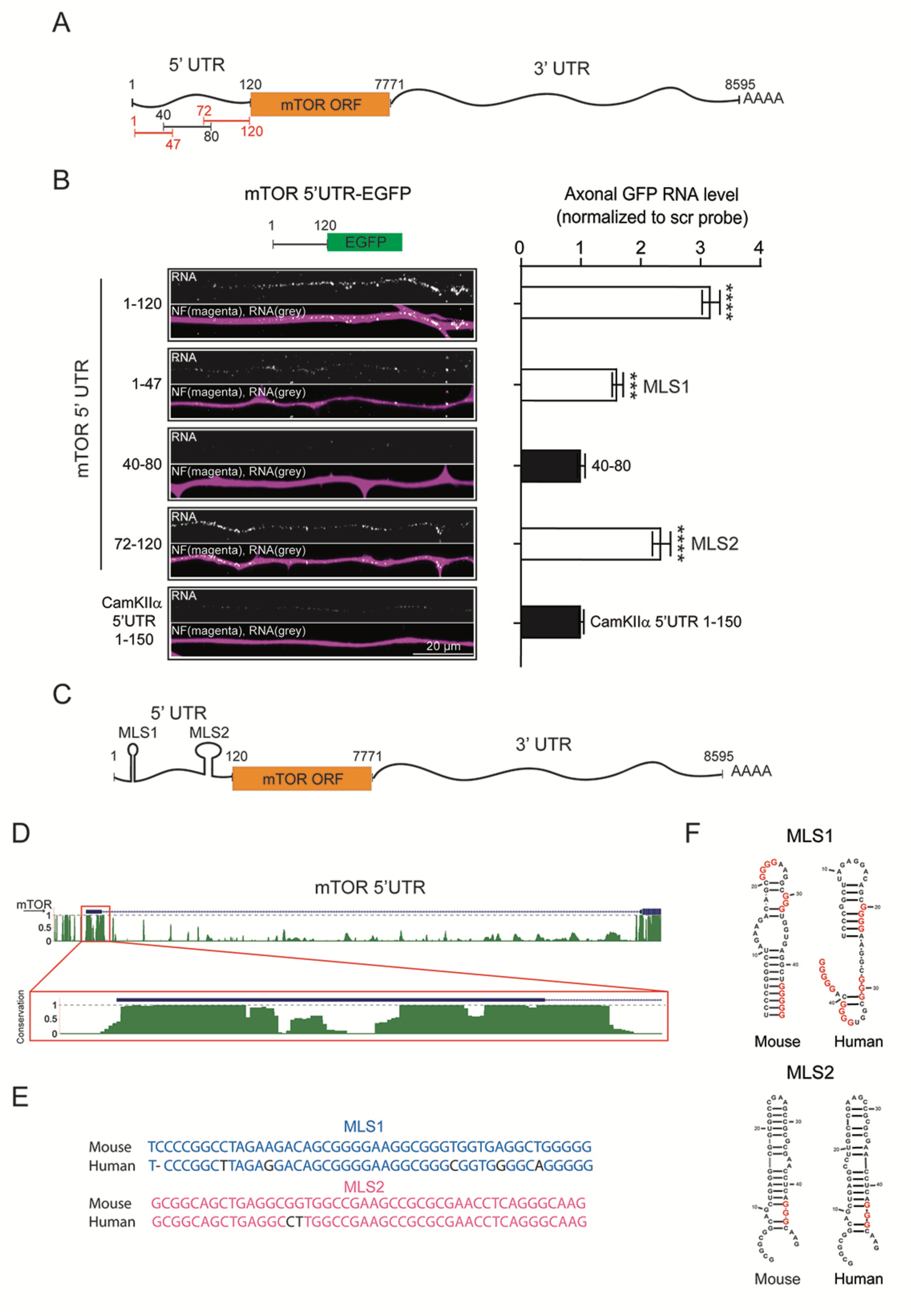
Two axonal localization signals in the mTOR 5’UTR. smFISH on cultured DRG sensory neurons transfected with constructs expressing EGFP reporter fused to sequence segments from mTOR 5’UTR. (**A**) Schematic of mTOR mRNA sequence showing the different segments tested. Axon-localizing 5’UTR segments highlighted in orange. (**B**) Representative axonal EGFP FISH images for 5’UTR fusion constructs, with quantifications as shown. CamkIIα 5’UTR 1-150 served as a negative control. Quantifications normalized to the scrambled (scr) control probe, n = 3 independent experiments, *** *p* < 0.001, **** *p* < 0.0001, one way ANOVA with Tukey’s post-test. (**C**) Schematic of mTOR mRNA showing positions of the delineated mTOR 5’UTR localization signals (MLS). (**D**) mTOR 5’UTR gene schematic (blue) from the UCSC genome browser, sequence conservation between 35 vertebrate species is shown underneath in green. Insert shows the mTOR 5’UTR first exon. (**E**) Sequence alignments of mouse versus human MLS1 and MLS2 from the mTOR 5’UTR, non-identical nucleotides in black. (**F**) Predicted secondary structures of mouse versus human MLS1 and MLS2.

The findings above indicate that two 5’UTR sites are involved in mTOR localization, in addition to the 3’UTR. We aimed to use gene editing with CRISPR/Cas9 to determine the relative contribution of the different MLS *in vivo* but first had to assess the likelihood of undesired effects upon mutation of the short 5’UTR. MLS2 is divided between the two exons of the 5’UTR by a 3.5 kbp intron, with 14 bases in the second exon (Extended Figure 1B). We therefore compared reporter RNA axon localizing activity for an exon 1 deletion construct retaining the 14 MLS2 nucleotides from exon 2 with a construct representing complete MLS1+MLS2 deletions and found that in both cases there was no EGFP localization to axons (Extended Figure 1C-E). Thus, it should be possible to avoid effects on mTOR gene structure by restricting editing to the first exon.

A second potential complication of mutating the mTOR 5’UTR is reported IRES activity within the MLS2-containing region [24]. To test whether deletion of exon 1 MLS2 nucleotides might disrupt mTOR IRES activity, we transfected HEK293 cells with EGFP cloned in fusion with mTOR 5’UTR sequences and tested effects of cap-dependent translation initiation inhibition on EGFP expression (Extended Figure 1F, G). The residual 14 nucleotides of mTOR exon 2 suffice to drive IRES-dependent reporter expression in contrast to a calreticulin UTR segment that lacks IRES activity [25], and likewise for mTOR 5’UTR sequence with deletion of exon 1 encoded MLS2 sequence (ι173-101). Thus, restricting gene editing to the first exon of mTOR should not affect IRES activity and should suffice to disrupt axon localization dependent on 5’UTR motifs.

### Deletion of both 5’UTR MLS generates mTOR hypomorphs

We applied CRISPR/Cas9 on heterozygotes for our previously described mTOR 3’UTR deletion line [16] to try to first generate gene-edited mice with small deletions or mutations within MLS1 and/or MLS2. However, multiple attempts to generate minimally mutated lines were not successful, and we routinely obtained mice with complete or almost-complete MLS1/2 deletions, with or without the mTOR 3’UTR deletion (ΔMLS1,2 and ΔMLS1,2/Δ3’UTR, Figure 2A). Both these lines were hypomorphs for mTOR expression, with reductions of 60-80% in both mRNA and protein levels for mTOR in brain and liver extracts, as well as in DRG neuron cultures (Figure 2B and Extended Figure 2A). There were no significant differences in mRNA degradation rates in actinomycin D-challenged DRG neuron cultures from the different lines (Extended Figure 2B, C), indicating that the reduced mTOR expression is not due to changes in RNA stability. Both mTOR hypomorph lines showed reduced body weight (Figure 2C) and non-Mendelian ratios for homozygous births with reduced litter sizes. Furthermore, we observed reduced brain sizes and weights in the hypomorph lines (Figure 2D). Thus, concomitant deletion of both 5’UTR MLS leads to mTOR hypo-morphism with effects on both body and organ size. We therefore turned to examination of individual 5’UTR MLS deletions, in combination with the 3’UTR.

**Figure 2.**
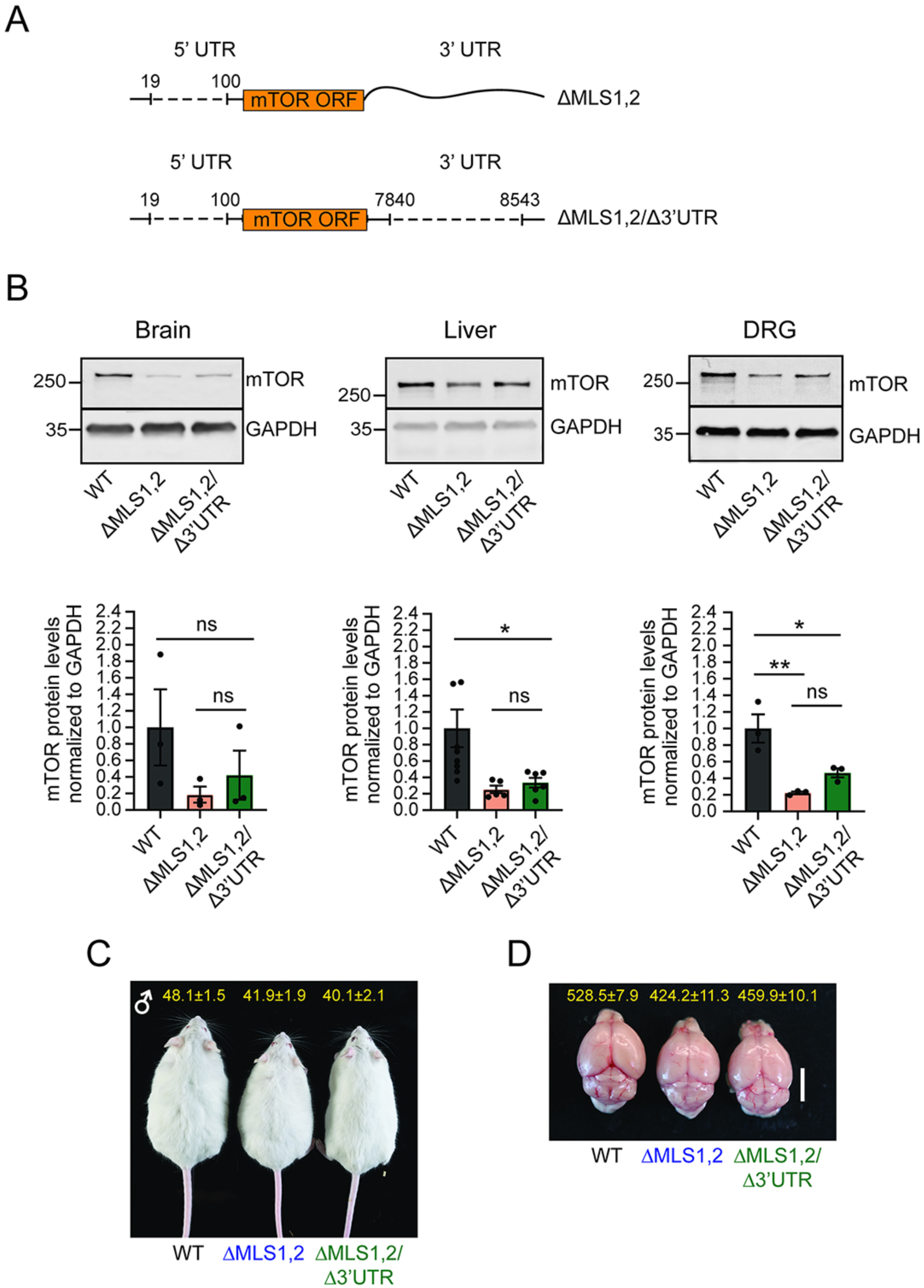
Combined deletions of both 5’UTR MLS generates mTOR hypomorphs with effects on animal and organ size. (**A**) Schematics showing deletions in the mTOR ΔMLS1,2 and mTOR ΔMLS1,2/Δ3’UTR mouse lines. (**B**) Representative Western blots showing mTOR protein levels in the indicated tissues of the mutant mice with quantifications below. (**C**) Representative images of wild-type and mutant 8-month-old male mice, with average + SEM body weights (gram) shown above. (**D**) Representative images of brains from 8-months old male mice, with average + SEM brain weights (mg) shown above, scale bar 5 mm.

### An MLS2 and 3’UTR deletion leads to subcellular mislocalization of mTOR

Further attempts to generate individual MLS1 or MLS2 mutants on 3’UTR mutant background generated a mouse line lacking MLS2 in the 5’UTR together with the 3’UTR deletion (Figure 3A). This mouse line, termed ΔMLS2/Δ3’UTR, retains most of the MLS1 sequence at the 5’ end of the transcript except for one nucleotide deletion. As with previous lines, there is no change in mRNA stability (Extended Figure 2B), but in contrast to the double-5’-MLS deletion lines the ΔMLS2/Δ3’UTR mice showed only modest reduction in mTOR mRNA levels (∼20%), with no significant change in mTOR protein levels in all tissues tested (Figure 3B and Extended Figure 3A). Moreover, and again unlike mice with deletion of both 5’UTR MLS, ΔMLS2/Δ3’UTR mice did not show changes in body weight (Extended Figure 3B, C).

**Figure 3.**
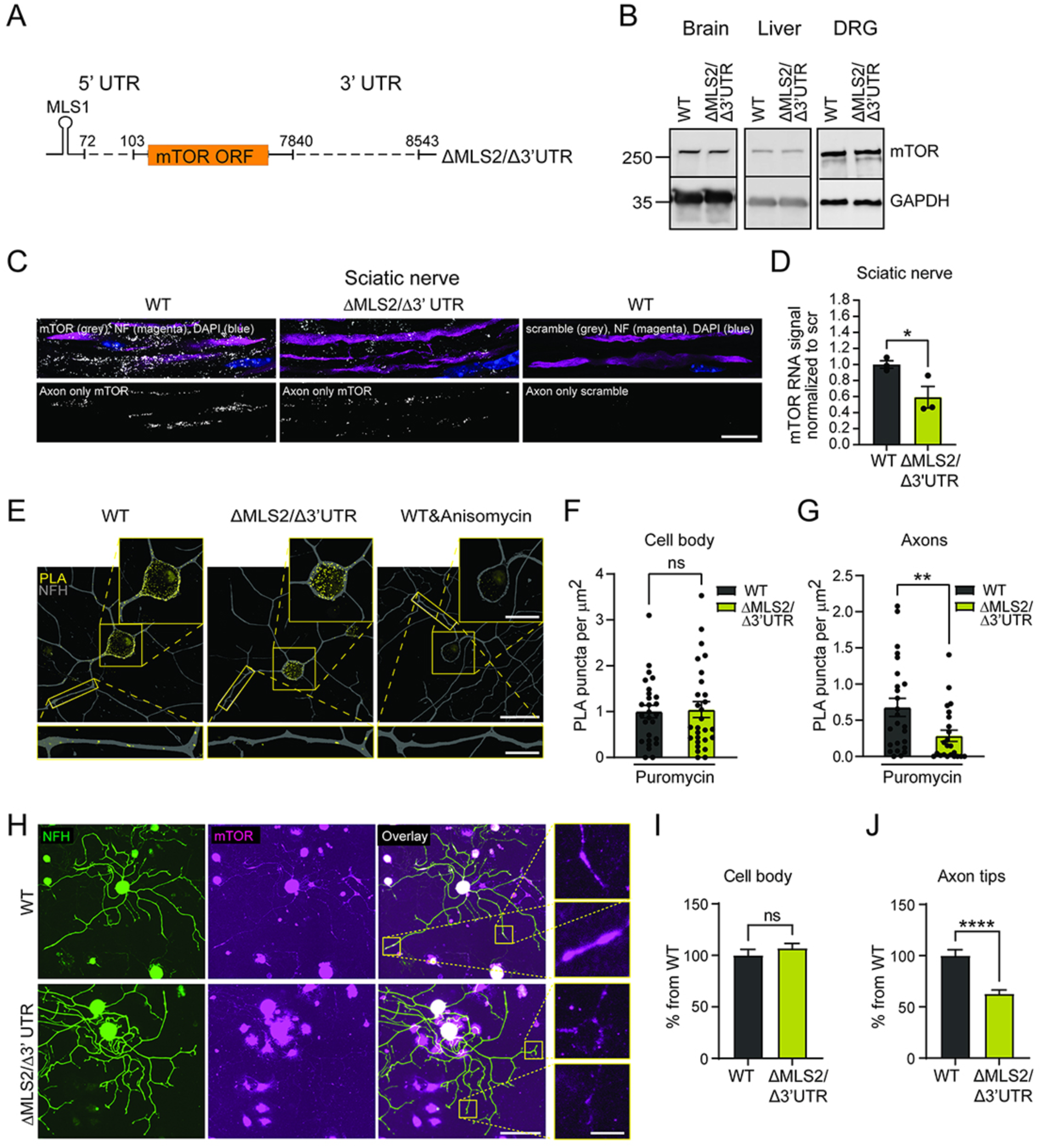
Mislocalization and reduced axonal levels of mTOR in the ΔMLS2/Δ3’UTR mouse line. (**A**) Schematic of the changes in the mTOR ΔMLS2/Δ3’UTR mouse line. (**B**) Western blots for mTOR in the indicated tissue extracts. (**C**) Representative, exposure-matched confocal images of smFISH (Stellaris) for mTOR mRNA or control scramble probe (grey) and neurofilament (magenta) immunostaining from sciatic nerve sections of wild-type (WT) and mTOR ΔMLS2/Δ3′UTR mice. Scale bar 10 µm. (**D**) Quantification of C, n = 3, means ± SEM, ns – nonsignificant, * p<0.05, unpaired t test. (**E**) Representative images of puromycin-PLA assay for mTOR on cultured DRG neurons. Scale bars: 50 µm (main image), 25 μm (Cell body ROI), 10 µm (axon ROIs). (**F**, **G**) Quantification of PLA puncta per µm^2^ in cell bodies (F) and axons (G), following normalization to a control anisomycin treatment. n=3; ** p <0.01, outliers removed by the ROUT method with Q=1% followed by unpaired two-tailed Mann-Whitney test. (**H**) Cultured DRG neurons immunostained for NFH (Green) and mTOR (magenta). Inserts (right) show axon tips. Scale bars 50 μm and 10 μm, respectively. (**I**, **J**) Quantification of mTOR protein levels in cell bodies (I) and axon tips (J), normalized to wild-type controls, n ≥ 35 cells from three independent cultures, **** p < 0.0001, t-test.

FISH analysis on sciatic nerve sections revealed a significant reduction in mTOR mRNA levels in sciatic nerve axons (Figure 3C, D). A puro-PLA assay [26, 27] showed approximately 50% reduction in local translation of mTOR in ΔMLS2/Δ3’UTR axons of cultured DRG neurons (Figure 3E-G). The apparent reduction in mTOR axonal translation was mirrored by a significant decrease in mTOR protein levels in axon tips of cultured sensory neurons from mTOR ΔMLS2/Δ3’UTR mutant mice (Figure 3H-J). Thus, combined deletion of MLS2 and the 3’UTR does not affect soma levels of mTOR, but mis-localizes axonal mTOR mRNA and causes a reduction in mTOR protein in axon tips.

### Attenuated axonal protein synthesis and increased axon growth in MLS2 and 3’UTR deletion neurons

Our previous work demonstrated that locally translated mTOR drives the translation of other proteins in axons [16]. We therefore compared local translation in cell bodies and axons of wild type versus mTOR ΔMLS2/Δ3’UTR mutant neurons after nerve injury. Mice were subjected to sciatic nerve crush, and neurons were cultured from L4/L5 DRG three days later (Figure 4A). DRG neurons from the non-injured contralateral side served as an additional comparison. Cultures were pulsed with puromycin to label nascent polypeptides, and new protein synthesis was quantified from puromycin immunostaining (Figure 4B, C). mTOR ΔMLS2/Δ3’UTR neurons show a significant reduction in puromycin labeling in axon tips, but not in cell bodies (Figure 4A-C), indicating that axonal deficits in mTOR specifically affect local translation in axons.

**Figure 4.**
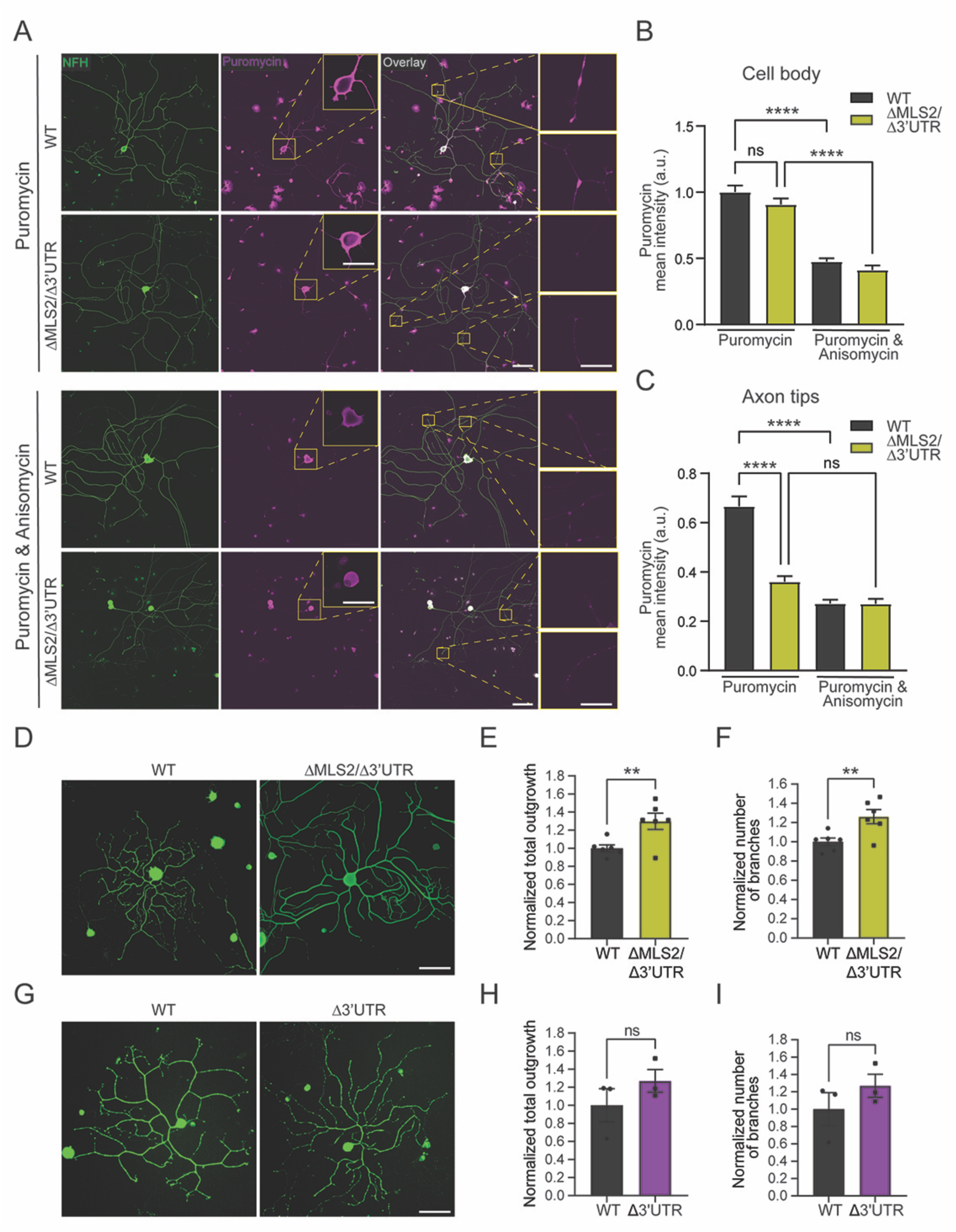
ΔMLS2/Δ3’UTR neurons show attenuated axonal translation and enhanced axonal growth. (**A**) Representative images of 24 hr cultures of L4/L5 DRG neurons from conditionally lesioned WT and ΔMLS2/Δ3’UTR mice, pulsed with puromycin for 10 min, fixed and immunostained for puromycin. Insets (right) show signal in axon tips. Scale bars: large image - 100 µm, cell body – 40 µm, axon tip 10 µm. (**B**, **C**) Quantification of puromycin signals in cell bodies (B) and axon tips (C), normalized to WT. n ≥ 25 cells from three independent cultures, **** p <0.0001, outliers removed by the ROUT method with Q=1% followed by Kruskal-Wallis test with Dunn’s post hoc. (**D**, **G**) Representative images of DRG neurons of the indicated genotypes 48 hr in culture immunostained for NFH. Scale bars: 100 µm. (**E, F**, **H**, **I**) Quantification of axonal outgrowth (E, H) and branching (F, I) of the indicated genotypes normalized to WT, n = 6 (ΔMLS2/Δ3’UTR) or n = 3 (Δ3’UTR), **p < 0.01, ns - not significant, Unpaired t-test.

Intrinsic neuronal growth is regulated by axonal transport of the RBP nucleolin and its mRNA cargos [12, 13], one of which is mTOR [16]. We therefore examined axonal growth of wild type, ΔMLS2/Δ3’UTR, and Δ3’UTR neurons, and saw significant enhancement of overall axonal growth and branching in ΔMLS2/Δ3’UTR neurons (Figure 4D-F). A similar trend was observed in Δ3’UTR neurons, but these did not reach statistical significance (Figure 4G-I). Thus, combined deletion of both 5’UTR and 3’UTR mTOR localization signals is required for significant effects on axon growth.

## Discussion

Where is mTOR and what is it doing there is a question posed by Betz and Hall over a decade ago [3]. Indeed, subcellular localization of mTOR is now appreciated as a critical determinant of its many activities, but the field has been mainly focused on how mTOR protein or mTOR signaling complexes are localized. RNA-driven localization of mTOR was demonstrated in both peripheral and central neurons, and was shown to affect diverse processes, including neuronal development [15, 17], synaptic communication [19], and axonal injury responses [16, 18]. Here we have shown that mRNA localization is encoded in both 5’ and 3’ mTOR UTRs, and that concomitant deletions in both UTRs are required to disrupt specific localization-dependent effects on neuronal growth. The existence of multiple non-overlapping MLS and their individual evolutionary conservation attests to the robustness and importance of mTOR mRNA localization.

RNA localization sequences have been most often reported in 3’UTRs [7, 28], giving rise to the notion that localization is encoded mainly in the 3’UTR, and translational control more in the 5’UTR [23, 29–33]. In this context, the occurrence of two independent localization motifs in the short 5’UTR of mTOR was unexpected. Nonetheless, a few other examples of 5’UTR localization motifs have been reported in the literature [20–22]. Neuritin (Nrn1) is similar to mTOR in that it has localization motifs in both 5’ and 3’UTRs [20], but in that case the 3’UTR motif suffices for axonal localization in cortical neurons [34], while the 5’UTR is necessary and sufficient for localization in sensory neurons [20]. The mTOR mRNA differs in that both 5’ and 3’UTRs can independently confer axonal localization in sensory neurons, likely conferring strong robustness for axonal localization of mTOR mRNA.

As noted, two independent mTOR MLS are encoded in the relatively short 5’UTR, and deletion of both caused marked reduction in mTOR expression levels. The reduced body sizes of these mTOR hypomorph lines are like those previously described for an mTOR hypomorph generated by insertion of a neomycin cassette to the mTOR gene, partially disrupting its transcription [35, 36]. Reduced mTOR expression in the 5’ tandem MLS deletion line is also likely due to reduced transcription, since we did not see changes in transcript stability, despite the significant reduction in transcript expression levels. Indeed, MLS1 deletion might affect transcription factor binding sites that were mapped within the 5’UTR [37]. Remarkably, both the 5’UTR double MLS mutant line and the previously described hypomorph line show that ∼75% reduction in mTOR expression is still compatible with life, at least in mice.

In contrast to the dual 5’UTR MLS deletion lines, deletion of only 5’UTR MLS2 together with the 3’UTR generated a true subcellular RNA localization deficit, without affecting overall mTOR protein expression. The specific reduction of axonal mTOR mRNA impacted nascent protein synthesis in axons without perturbing synthesis in cell bodies and led to enhancement of axonal growth. This effect did not reach statistical significance in the mutant with 3’UTR-only deletion, and indeed our previous study found that the main neuronal phenotype in that line was reduced sensory neuron survival after axonal injury [16]. Previous studies have shown that sequestration of nucleolin and its cargo mRNA KPNB1 away from axons and their concentration in cell bodies likewise accelerate axonal growth [11–14] and regulate presynapse composition [38]. Our current findings support mTOR involvement in this axonuclear communication mechanism for regulating cell size and neuron growth [11, 39]. Thus, subcellular localization of mTOR is ensured by multiple RNA localization motifs to regulate neuron growth.

## Methods

### Animals

All animal procedures were approved by the Weizmann Institute and University of South Carolina Institutional Animal Care and Use Committees (IACUC). Adult mice of 2-8 months age and male Sprague Dawley rats (175–250 g) were used and had access to food and water ad libitum.

### Cloning mTOR 5’UTR sequences fused to EGFP

The mTOR UTR sequences were amplified from purified tail DNA of wild type ICR mice, and were cloned into a modified pEGFP-C2 plasmid vector using restriction free cloning [40]. All the EGFP constructs were under a CMV promotor. The rat CamkIIα 5’UTR (NCBI, XM_039096593.1) was used as a non-localizing control.

### DRG neuron culture

DRG neuron culture was done as previously reported [13, 27]. Briefly, DRG were dissected from adult mice (2-8 months old) and collected in 1 x Hanks’ buffered saline solution (HBSS), followed by two steps of enzymatic digestions of connective tissues with 20 u/ml papain (Sigma #P-4762) for 20 min and a mixture of 1.6 mg/ml Collagenase II (Worthington # CLS-2) and 2.4 mg/ml Dispase II (Roche #165859) for another 20 min. The enzymatic digestion was followed by mechanical dissociation using a fire polished 1 mm glass pasteur pipette and neurons enriched by centrifugation at low speed (1000 x g) through a 20% percoll cushion (Sigma #P1644) in L-15 media (Sigma #L-5520). Neurons were washed and suspended in F-12 medium (GibcoBRL #21765-029) supplemented with 20 µM Ara-C (Sigma #C6645) and seeded on glass coverslips pre-coated with 0.01% poly-L-lysine solution (sigma P-4832) for at least 16 hr at 4°C, followed by 8 µg/ml laminin (Invitrogen, 23017-015) for at least 2.5 hr at 4°C.

### Fluorescence in situ hybridization (FISH)

FISH experiments were done as described previously [16]. Briefly, adult rat DRG neurons were transfected immediately after dissociation with the EGFP constructs by electroporation using the Amaxa nucleofector II (Lonza), the neuron nucleofector VGP-1003 kit Rat (Lonza), and program G-013. The cells were incubated in culture for 36 hr then fixed with 2% PFA, permeabilized with 0.3% triton x100. Oligonucleotides corresponding to the EGFP coding sequence (#NCBI accession #U55762.1, residues 172-222 and 531-581) modified with 5′-amino C6 at four thymidines per oligonucleotide (IDT, Inc.) were covalently labeled with digixogen (DIG) using digoxigenin succinamide ester (Roche) and used for hybridization. Scrambled probes were used as control for background labeling. The probe hybridization was done in hybridization buffer (2% BSA, 1mM Ribonucleoside-vanadyl complex, 20mM NaBO_4_ in 1x SSC buffer) containing 1.66 ng/µl probe and incubated at 37°C for 5-10hr. Probe detection and immunofluorescence were done in blocking buffer (50 mM Tris pH 7.5, 1% BSA Heat shock BSA, 1% Protease free BSA) using mouse anti DIG, and Chicken anti NF. Secondary antibodies were Cy3 conjugated donkey anti mouse (1:200, Jackson ImmunoRes.) and Cy5 conjugated donkey anti chicken (1:200, Jackson ImmunoRes.). Images were obtained by a Leica DMI6000 epifluorescent microscope with ORCA Flash ER CCD camera (Hamamatsu). Image J was used to determine mRNA and protein levels in axons and cell bodies.

FISH on tissue sections was conducted as previously described [16], with minor modifications. Briefly, sciatic nerves were excised from WT and mTOR ΔMLS2\Δ3’UTR mice, fixed in 2% PFA for 6 hr at 4°C, washed in 1x PBS and incubated in 30% sucrose solution for at least 3 days at 4°C, before embedding in O.C.T (Tissue Trek, 4583). 20 µm thick longitudinal sections were collected using a cryostat microtome instrument and stored at -20°C until used. Sections were brought to room temperature, washed three times with 1 x PBS, and incubated with 20 mM glycine in 1 x PBS for 10 mins followed by 0.25 M NaBH_4_ in 1 x PBS for 10 mins to quench autofluorescence. Sections were then washed with 0.2 M HCl for 10 min, permeabilized with 1% Triton-X100 in 1 x PBS for 2 min then washed shortly with 1 x PBS following by equilibration in 2 x SSC with 10% formamide for 10 min. Fluorescence probe hybridization was done in hybridization buffer (2 x SSC, 10% formamide, 10% Dextran sulfate, 1 mg/ml E.coli tRNA, 2mM RVC, 200 µg/ml BSA) containing fluorescently labeled ’Stellaris’ probes (1:100, Biosearch Tech.) and mouse anti-NF (Novus, NBP2-29435, 1:200) and placed in a humidity chamber for overnight incubation at 37°C. Washes of the probe were done in 2 x SSC with 10% formamide at 37°C for 30 min followed by two washes in 2 x SSC for 5 min and then equilibration in 1% Triton-X100 in PBS for 5 min before incubation with donkey anti-mouse FITC antibody (1:200) in 0.3% Triton-X100 + 10X blocking buffer (1:100; Roche) for 1 hr at room temperature. The secondary antibody was washed with 1 x PBS for 5 min and the sections were fixed again with 2% PFA for 15 min and then washed in 1 x PBS for 5 min, rinsed in DEPC-treated water, and mounted using Prolong Gold Antifade (Invitrogen).

FISH/IF images for mTOR mRNA and NF protein co-localization were collected by confocal microscopy using a Leica SP8X confocal microscope with HyD detectors. Scramble probes were used to set image acquisition parameters that would not acquire any nonspecific signal. The Z stacks images were taken of distal axons separated from glial cells with 63X oil-immersion objective (1.4 NA). Z stacks were post-processed by Huygen’s deconvolution using the HyVolution module of Leica LASX software. The colocalizing signals were extracted in single optical planes using an ImageJ plugin (https://imagej.net/Colocalization_Analysis).

### mTOR sequence alignment and secondary structure prediction

mTOR sequence conservation in 35 vertebrate species was assessed in the mouse genome browser (GRCm39/mm39) using PhastCons. Mouse mTOR UTR sequences were obtained from the wild type Hsd:ICR (CD-1) mice and aligned to human mTOR sequence (NM_001386500.1) by global multiple sequence alignment using Clustal Omega[41] and visualized by Jalview software version 2.11.4.0 [42]. Secondary structure prediction was done in Mfold [43] and visualized with RNAcanvas [44] webservers.

### SDS-PAGE and Western blot

Protein samples were run on 4-15% acrylamide gradient gels (Bio-Rad) at a constant 120V for 1 hr and then transferred to a nitrocellose membrane using a Trans-blot Turbo system (Bio-Rad). The membrane was blocked in 5% milk solution in 1 x TBST buffer for 1 hr at room temperature, followed by incubation with the primary antibody in 1 x TBST overnight at 4°C. Following three 15 min washes with 1 x TBST, the membrane was incubated for 45 min at room temperature with the secondary antibody (goat anti-mouse HRP (1:10,000) or goat anti-rabbit HRP (1:10,000), Bio-Rad cat. 1706516 and 1706515, respectively). The membrane was washed again in 1 x TBST for three times 15 min and chemiluminescence was assayed using the Radiance Q kit (Azure Biosystems AC2101) in an ECL scanner (Amersham Imager 680).

Antibodies used were as follows:

| Antibody | dilution | Company and catalog # |
| --- | --- | --- |
| Rabbit anti Nucleolin | 1:1,000 | Cell Signaling cat. D4C70 |
| Mouse anti puromycin | 1:1,000 | Merck, cat. MAB343 |
| Rabbit anti puromycin | 1:1,000 | ABclonal, Cat. A23031 |
| Rabbit anti EGFP | 1:1,000 | Abcam, cat. ab6556 |
| Rabbit anti mTOR (monoclonal) | 1:100/1,000 IF/WB | Cell Signaling cat. 2983S |
| Mouse anti GAPDH | 1:50,000 | Merck, cat. MAB374 |
| Mouse anti actin (C4) | 1:1,000 | MP Biomedicals, cat. 08691001 |
| Chicken anti NFH | 1:1,000 | Abcam, cat. ab72996 |
| Mouse anti NF | 1:200 | Novus, cat. NBP2-29435 |
| Goat anti mouse HRP | 1:10,000 | Bio-Rad, cat. 1706516 |
| Goat anti rabbit HRP | 1:10,000 | Bio-Rad, cat. 1706515 |
| Donkey anti chicken Alexa 488 | 1:1,000 | Jackson Immuno Research cat. 703-545-155 |
| Donkey anti rabbit Alexa 647 | 1:1,000 | Jackson Immuno Research cat. 711-605-152 |
| Donkey anti mouse Alexa 647 | 1:1,000 | Jackson Immuno Research cat. 715-605-151 |
| Mouse anti DIG | 1:500 | Jackson immuno-research cat. 200-002-156 |

### Mouse gene editing using CRISPR/Cas9

gRNAs were designed using a combination of in-silico design tools, including the MIT CRISPR design tool [45] and sgRNA Designer, Rule set 2 [46], in the Benchling implementations (www.benchling.com), SSC [47] and CRISPOR [48]. Two gRNAs were designed to target Exon 1 of mTOR (see Oligonucleotide list below). A single-stranded oligodeoxynucleotide (ssODN) donor repair template containing mTOR 5’UTR 47-80 sequences flanked asymmetrically by homology arms to each of the 5′ and 3′ insertion sites was also designed, as shown in the Oligonucleotide list below. Cas9 nuclease, crRNA, tracrRNAs and ssODN were purchased from Integrated DNA Technologies (IDT). mTOR mutant mice were generated in the gene editing core facility at the Weizmann Institute of Science using CRISPR/Cas9 genome editing in isolated one-cell mouse embryos as described [49]. mTOR Δ3’UTR mice homozygous males [16] were mated with WT Hsd:ICR (CD-1) females that were purchased from Envigo, Israel and maintained in specific pathogen-free (SPF) conditions. Mice were maintained on a 12 hr light/dark cycle, and food and water were provided ad libitum. Cas9-gRNA ribonucleoprotein (RNP) complexes together with donor repair template were delivered to one-cell embryos via electroporation, using a Biorad Genepulser. Electroporated embryos were transferred into the oviducts of pseudopregnant ICR females (Envigo, Israel). Genomic DNA from F0 pups was analyzed at weaning by PCR and Sanger sequencing using primers listed below. Mice carrying the selected modifications were bred to wild-type Hsd:ICR (CD-1) mice to determine germline transmission. Mice were backcrossed for at least three generations with wild-type Hsd:ICR (CD-1) to minimize off-target effects, including at least one male-mediated cross for the Y chromosome.

### List of Oligonucleotides (ODN)

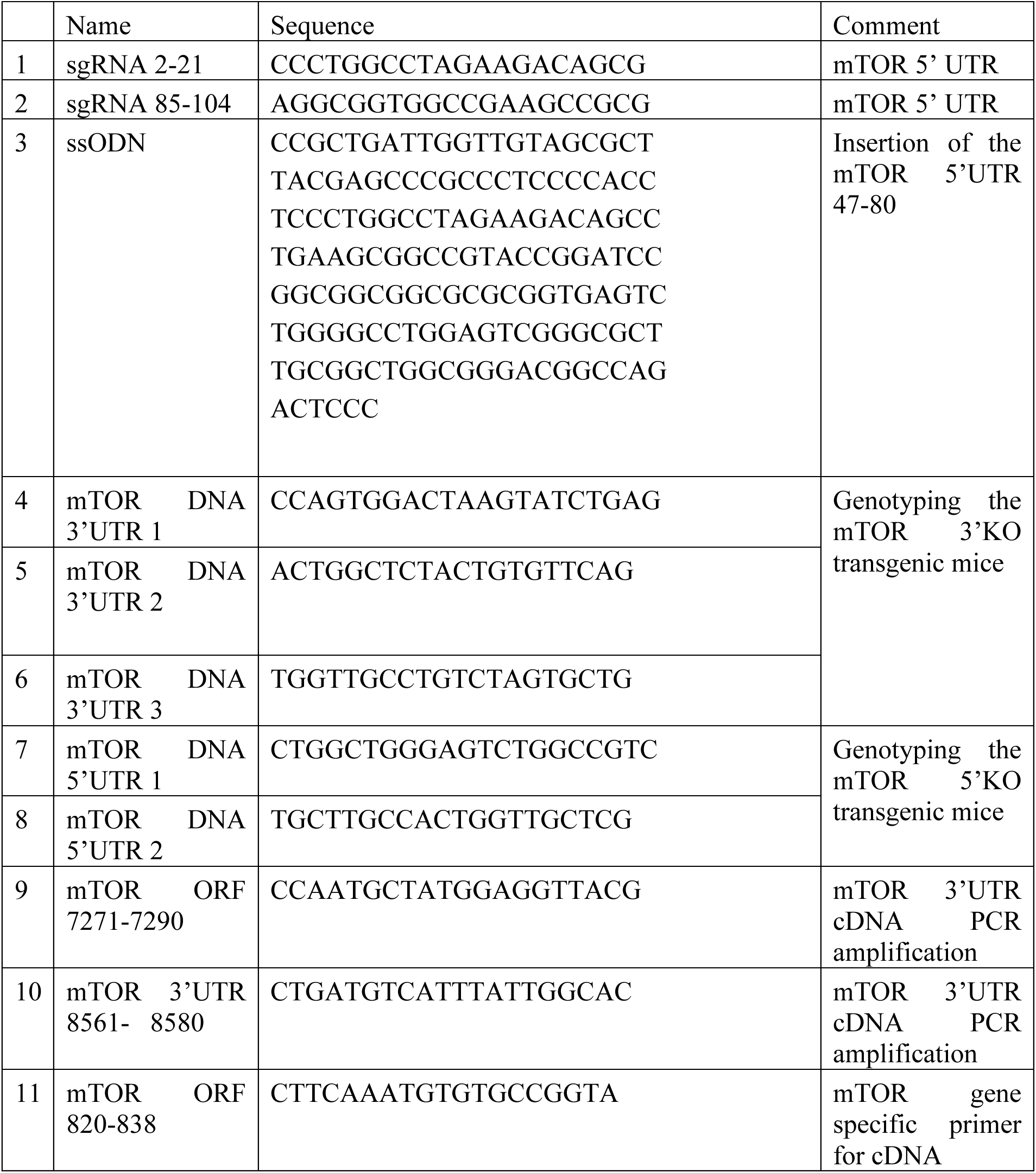
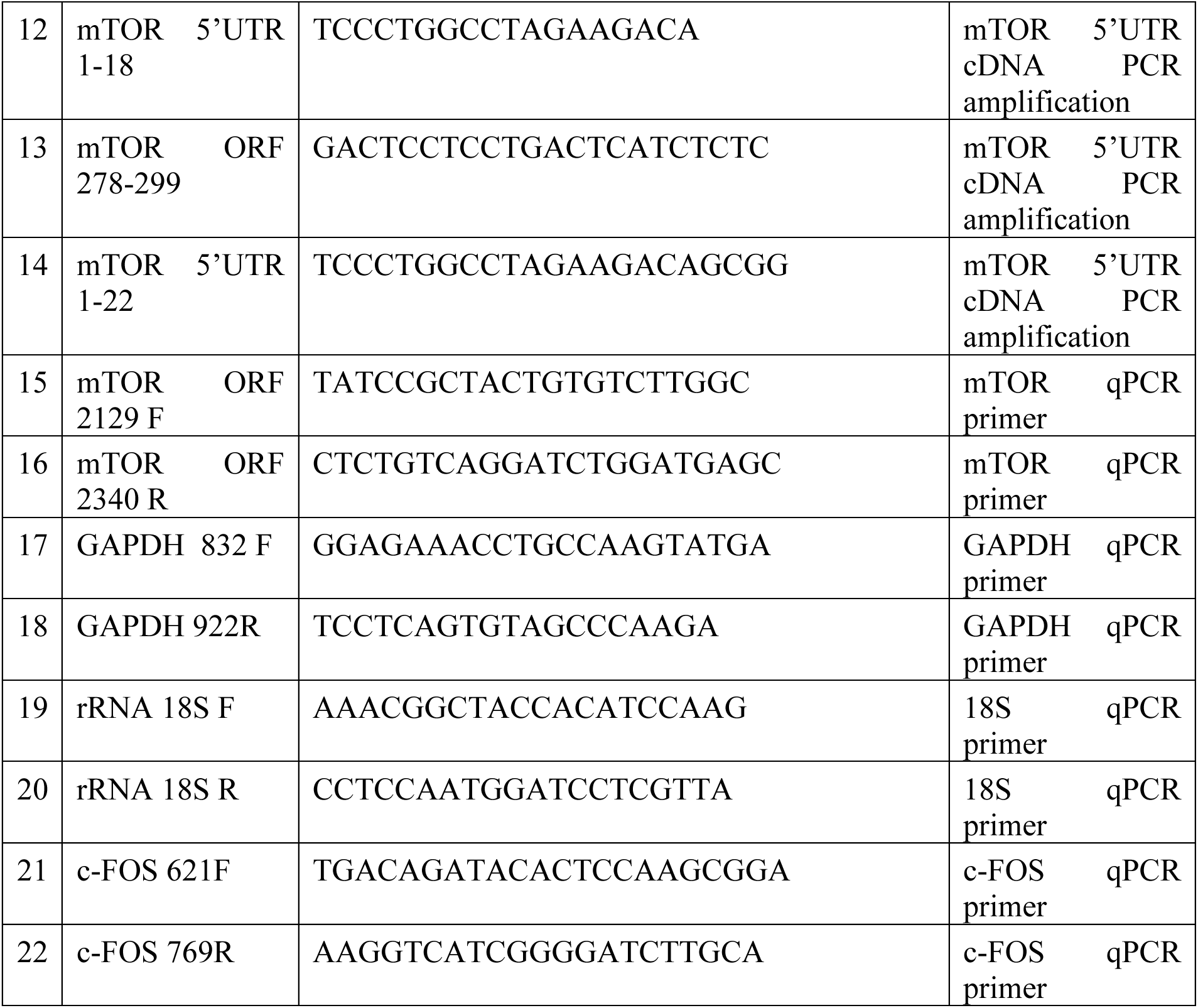

### Genotyping mTOR UTR edited mice

Ear clips were incubated in DNA extraction buffer (10 mM Tris pH 8.0, 100 mM NaCl, 10 mM EDTA, 0.5% SDS) supplemented with 0.4 µg/µl proteinase K at 55°C for at least 6 hr followed by 45 min of inactivation at 85°C. DNA lysates were cooled to 4°C, and PCR reactions were done with Taq DNA Polymerase 2x Master Mix RED (Ampliqon cat. 5200300), 1 µM of each primer and 1 µl DNA lysate. For the mTOR 3’ UTR, three primers were used in a single reaction (#4-6 in the ODN list). PCR reaction: 95°C for 5 min, 95°C for 30 s, 65°C for 30 s, 72°C for 30 s, 33 cycles followed by 10 min at 72°C.

### RNA extraction and quantification

RNA was extracted from the brain and liver using the RNeasy mini kit (Qiagen). The tissues were suspended in RLT buffer (Qiagen) and homogenized using a plastic pestle, followed by centrifugation at 10,000 x g for 5 min at 4°C to remove tissue debris. Supernatants were loaded on the RNeasy mini column following manufacturer’s instructions. For the RNA stability test, DRG neurons were grown for 24 hr and transcription was inhibited by adding 5 µM actinomycin D (Sigma cat. A1410) to the growth medium. RNA was extracted at the indicated time points by the RNeasy micro kit (Qiagen) and the RNA concentrations were measured using the Qbit 2.0 flourometer (Invitrogen). Reverse transcriptase (RT-PCR) reaction was done with equal amounts of RNA taken from all samples for the generation of cDNA using the superscript III kit (Invitrogen cat. 18080051) with random hexamers according to manufacturer’s instructions. Quantitative PCR (qPCR) was carried out using Perfecta Sybrgreen master mix (Quanta cat. 95073-012-2) and 0.5 µM primers concentration in the VIIA 7 real time PCR system (Invitrogen).

### IRES activity test

Human Embryonic Kidney (HEK) 293 cells were grown in DMEM media (Thermo Fisher, cat. 41965-039) supplemented with 10% fetal bovine serum (FBS) and 1 mM sodium pyruvate. Upon reaching ∼70% confluency, cells were transfected with 1 µg of each EGFP plasmid, in duplicates, using the jetPEI kit (Polyplus, cat. 101-10N) according to manufacturer’s instructions. After 4 hr, cells were incubated with either 100 µM 4EGI or equal volume of DMSO (vehicle) for an additional 6 hr. Cells were washed once with 1 x PBS, and proteins were extracted with RIPA buffer (150 mM NaCl, 1% Triton x-100, 0.5% Na-deoxycholate, 50 mM Tris pH 8.0, 0.1% SDS) supplemented with protease inhibitor (Roche cat. 11873580001), the lysate was incubated at 4°C for 20 min, sonicated and centrifuged at 10,000 x g for 10 min in 4°C. Supernatant was taken to read protein concentration with BCA kit (ThermoFisher, cat. 23225) and 20 µg proteins were loaded on polyacrylamide gel and subjected to Western blot as described above with rabbit anti EGFP (1:1,000) and mouse anti Actin (1:1,000) antibodies.

### Sciatic nerve injury and DRG puromycinylation

Sciatic nerve injury and puromycin labeling of DRG neurons in culture were done as previously described [16]. In brief, mice were anesthetized with ketamine\xylazine, the sciatic nerve was exposed by a small incision through the skin and muscle above mid-thigh and then crushed two times for 30 seconds each at the same position but at different angles for maximal crush effectiveness. The incision area was then closed using sterile wound clips. Only one sciatic nerve was injured per animal while the contralateral side was used as a naive control. Mice were sacrificed three days following injury, and L4/5 DRG neurons were grown in culture for 48 hr as described above. After 48 hr, the DRG were pre-incubated with 40 µM anisomycin (Sigma cat. A9789) for 1 hr followed by incubation with 5 µM puromycin (Sigma cat. P8833) for 10 min in culture media. After fixation (4% PFA in 1 x PBS for 20 min) cells were washed in 1 x PBS and permeabilized (5% normal donkey serum, 1 mg/ml BSA, 0.1% triton x 100 in 1 x PBS) for 45 min. The cells were then incubated with the following primary antibodies: chicken anti NFH, mouse anti puromycin or rabbit anti mTOR and in blocking buffer (5% normal donkey serum, 1 mg/ml BSA in 1 x PBS) overnight at 4°C. Cells were washed in 1 x PBS, and secondary antibodies were added for 45 min at room temperature. Following thorough washes, cells were mounted on glass slides using flouromount-G (Southern Biotech cat. 0100-01). Image acquisition for cell bodies, growth cones and axon tips was on a Nikon Ti-LAPP illumination system using an Andor EMCCD camera. The images were captured at X40 magnification with long distance Nikon super plan fluor ELWD objective (NA 0.6) and analyzed by NIS-ELEMENTS software.

### Axonal outgrowth and immunofluorescence

DRG neurons were grown in culture for 48 hr, fixed for 20 min in 4% PFA in 1 x PBS at room temperature, and permeabilized with 5% normal donkey serum, 1 mg/ml BSA, 0.1% triton x 100 in 1 x PBS for 45 min. Staining with the first antibody was done overnight at 4°C (mTOR, 1:100) or for 1 hr at room temperature (NFH 1:1,000). For axonal outgrowth experiments, images were taken on an ImageXpress Micro (Molecular Devices) system using a 10x objective and analyzed with MetaXpress software (Molecular Devices), using the neurite length package. Neurons were included in analyses only if the maximal process length was at least twice the cell body diameter. Acquisition of images for cell bodies, growth cones and axon tips were done on a Nikon Ti-LAPP illumination system using an Andor EMCCD camera. The images were captured at X40 magnification with long distance Nikon super plan fluor ELWD objective (NA 0.6) and analyzed by NIS-ELEMENTS software.

### Proximity ligation assay (PLA)

Proximity ligation assays were done as previously described [26]. Briefly, DRG neurons were grown in culture on 13 mm coverslips for 48 hr and puromycin labeling, fixation and permeabilization was done as described above. The cells were then incubated with rabbit anti-mTOR (Cell Signaling cat. 2983S, 1:100) and mouse anti-puromycin (1:5,000) antibodies in blocking buffer overnight at 4°C. The proximity ligation assay was done using the Duolink kit (Sigma: PLA probe anti-mouse minus DUO92004, anti-rabbit plus DUO92002, with detection using Far-Red DUO92013) according to the manufacturer’s protocol. Following the PLA assay, cells were washed in 1 x PBS, incubated with chicken anti-NFH (1:2,000) antibody for 45 min at room temperature. After another wash in 1 x PBS, secondary antibody donkey anti chicken Alexa Fluor 488 (Jackson immunoresearch 1:1,000) and DAPI (1:10,000) were added for 45 min at room temperature. The coverslips were mounted with flouromount-G (Southern Biotech cat. 0100-01) and images were captured on an Olympus fluoview FV10i confocal microscope at X60 magnification using an UPLSAP60xW objective (NA 1.2), followed by analysis with ImageJ. NFH staining was used to generate a mask for the quantification of PLA puncta. The cell body area was manually selected using the NFH channel and PLA puncta were counted using the Analyze particle function. For axonal quantification, the cell body mask was removed to generate axon only mask and PLA puncta in the axons were quantified as above. The number of PLA puncta was normalized to the mask area.

### Protein extraction from mouse organs

Proteins were extracted from adult mouse brain and liver and immediately frozen in liquid nitrogen until further analysis. For DRG, neurons were grown in culture as described above for 48 hr. Tissue was suspended in RIPA buffer (150 mM NaCl, 1% Triton x-100, 0.5% Na-deoxycholate, 50 mM Tris pH 8.0, 0.1% SDS) supplemented with protease inhibitor (Roche cat. 11873580001) and homogenized using a plastic pestle in an Eppendorf tube on ice. The lysate was sonicated (3 x 10 s with 10 s) and cleared of cell debris by centrifugation (10,000 x g, 4°C). Protein concentrations of supernatants were measured by BCA assay (Thermo Fisher cat. 23225). 20 µg of protein from each sample was used for Western blot analysis using the rabbit anti mTOR (1:1,000) and the mouse anti GAPDH (1:5,000).

### Statistical methods

Data are shown as mean ± standard error of the mean (SEM), unless otherwise noted. Groupwise analyses were conducted by one- or two-way ANOVA with post hoc tests as described in the figure legends. A Chi-squared test was conducted for Mendelian genotype distributions against the expected Mendelian ratio. Pairwise analyses were conducted by either two-tailed unpaired Student’s t tests, non-parametric Mann–Whitney U test or Kolmogorov-Smirnov test. Where indicated, outliers were removed by the ROUT method with Q=1% followed by unpaired two-tailed Mann-Whitney test. All statistical analyses were conducted using either GraphPad Prism or R software. Statistically significant p values are shown as * p < 0.05, ** p < 0.01, *** p < 0.001 and **** p < 0.0001.

## Acknowledgments

We thank Elina Berkovitz, Golda Damari and Sima Peretz of the WIS Transgenic Facility for their great assistance in generating the CRISPR mouse models, and Dalia Gordon for helpful comments and criticism throughout the project.

## Funding

We thank the following funding sources - Dr. Miriam and Sheldon G. Adelson Medical Research Foundation (MF, JLT), European Research Council Advanced Grant GrowthSINE 101141522 (MF), National Institutes of Health R01NS117821 (JLT), South Carolina Spinal Cord Injury Research Fund (PKS, JLT), Merkin Peripheral Neuropathy and Nerve Regeneration Center (PKS), Weizmann Center for Research on Injury and Regeneration (MF), Sagol Weizmann-MIT Bridge Program (MF), The Company of Biologists travel grant (NS), the Chaya Professorial Chair in Molecular Neuroscience (MF), and the South Carolina Smartstate Chair in Childhood Neurotherapeutics (JLT).

---

**Extended Figure 1.**
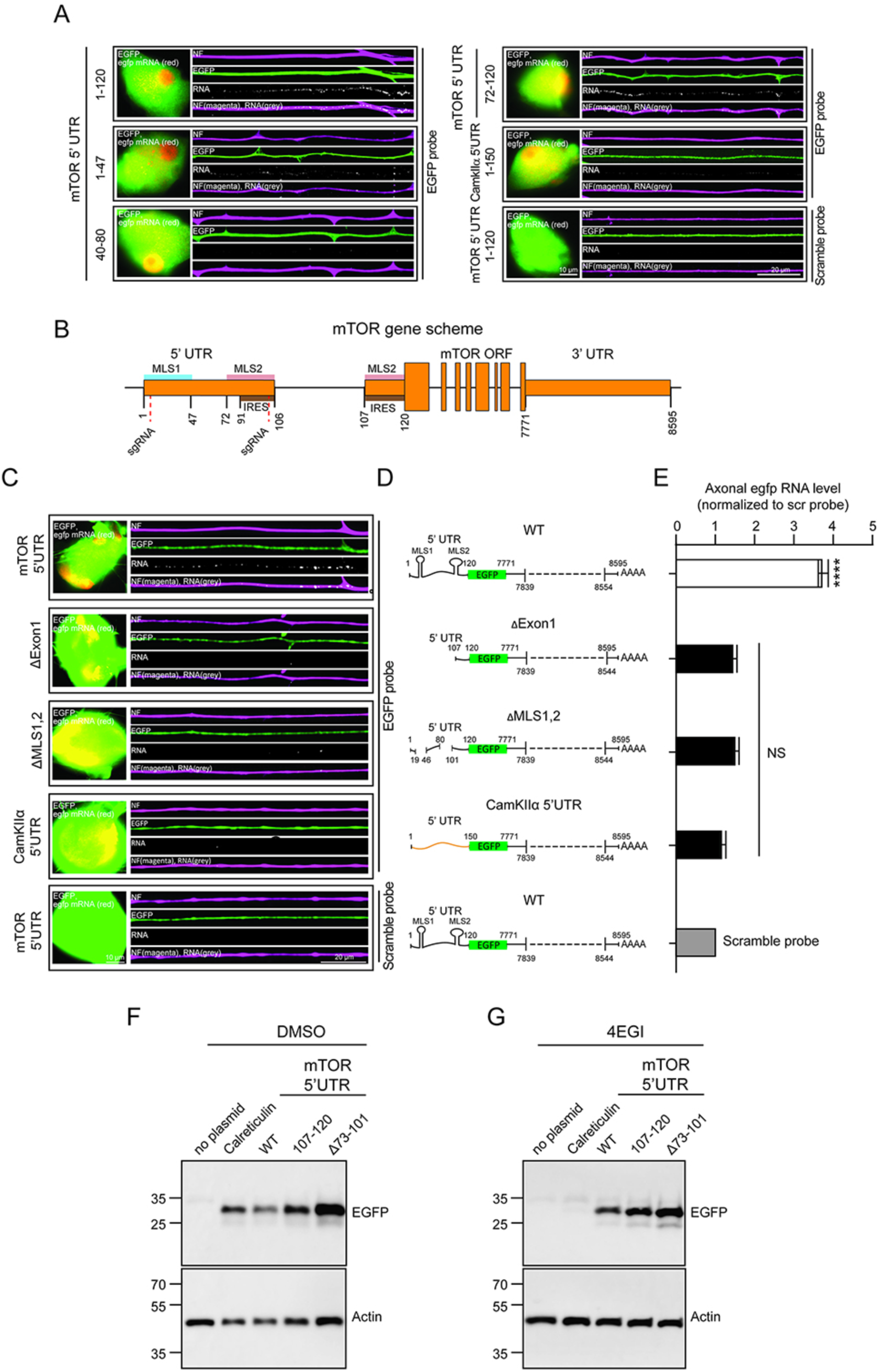
Discriminating RNA localization and IRES activities in the mTOR 5’UTR. (**A**) EGFP coding sequence was cloned together with mTOR UTR segments or control sequences as indicated and transfected to DRG neurons in culture. Representative epifluorescent images of immunofluorescence for EFGP protein in green, and smFISH for egfp mRNA in cell bodies (red) and axons (grey) with neurofilament (NF, magenta) as neuronal marker. CamKIIα 5’UTR 1-150 was used as a negative control. Scale bars 10 μm for somata, 20 μm for axons. (**B**) Schematic of mTOR gene structure showing locations of the two 5’UTR MLS, a previously reported 5’UTR segment containing IRES activity [24], and intron/exon boundaries for the UTRs. Numbers below the scheme indicate position along the mTOR mRNA. (**C**-**E**) Axon localization of EGFP reporter RNA in sensory neurons after transfection with the indicated mTOR UTR fusion constructs (5’ UTR 1-120, ΔExon1 leaving 5’UTR 107-120, ΔMLS1,2 leaving 5’UTR 1-18, 47-79, 102-120) or CamKIIα 5’UTR as control. (**C**) Representative FISH images of cell bodies and axons transfected with the constructs indicated in **D**. (**E**) Quantification of axonal egfp RNA levels, n = 3, **** *p* < 0.0001, one way ANOVA with Dunnett’s post-test. (**F**, **G**) Representative Western blots for EGFP and actin from HEK293 cell extracts transfected and incubated as shown. Calreticulin 5’UTR served as a control for IRES activity in these experiments. Cells were incubated with DMSO or eIF4E/eIF4G inhibitor (4EGI-1 at 100 µM) for 6 hr in culture before extraction.

**Extended Figure 2.**
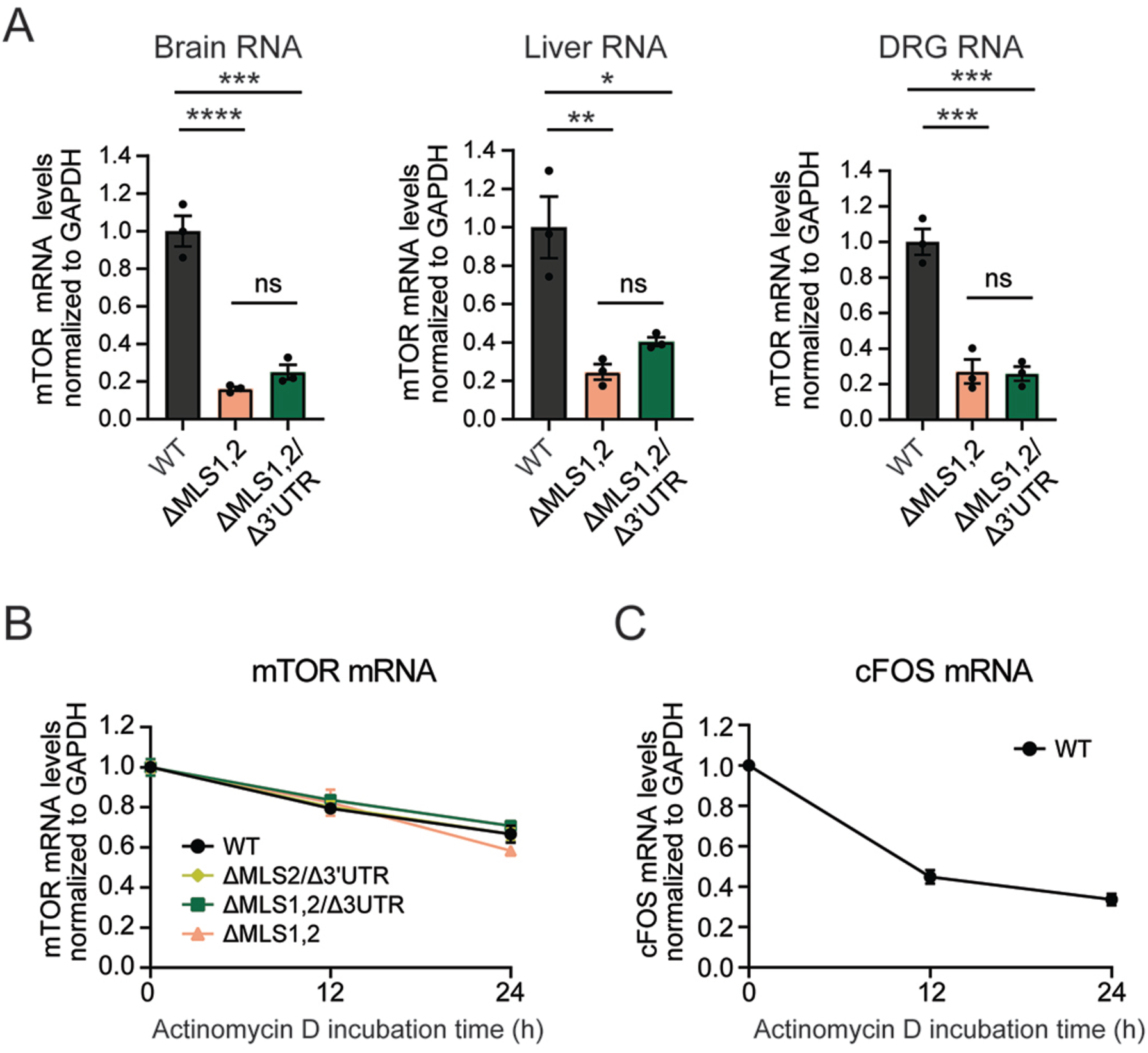
MLS1/2 combined deletion mouse lines express reduced levels of mTOR but retain mTOR mRNA stability. (**A**) mTOR mRNA levels quantified by qPCR in brain, liver and DRG from the indicated mouse lines. n ≥ 3, * *p* < 0.05, ** *p* < 0.01, *** *p* < 0.001, and **** *p* < 0.0001. One-way ANOVA with Tukey’s post hoc test. (**B, C**). Cultured DRG neurons were incubated with vehicle or the transcription inhibitor actinomycin D (5 μM) for 0, 12 and 24 hr. mTOR (B) and cFOS (C) mRNA levels were quantified by RT-qPCR. n = 3, p < 0.05 between time points, two-way ANOVA with Sidak’s post-test.

**Extended Figure 3.**
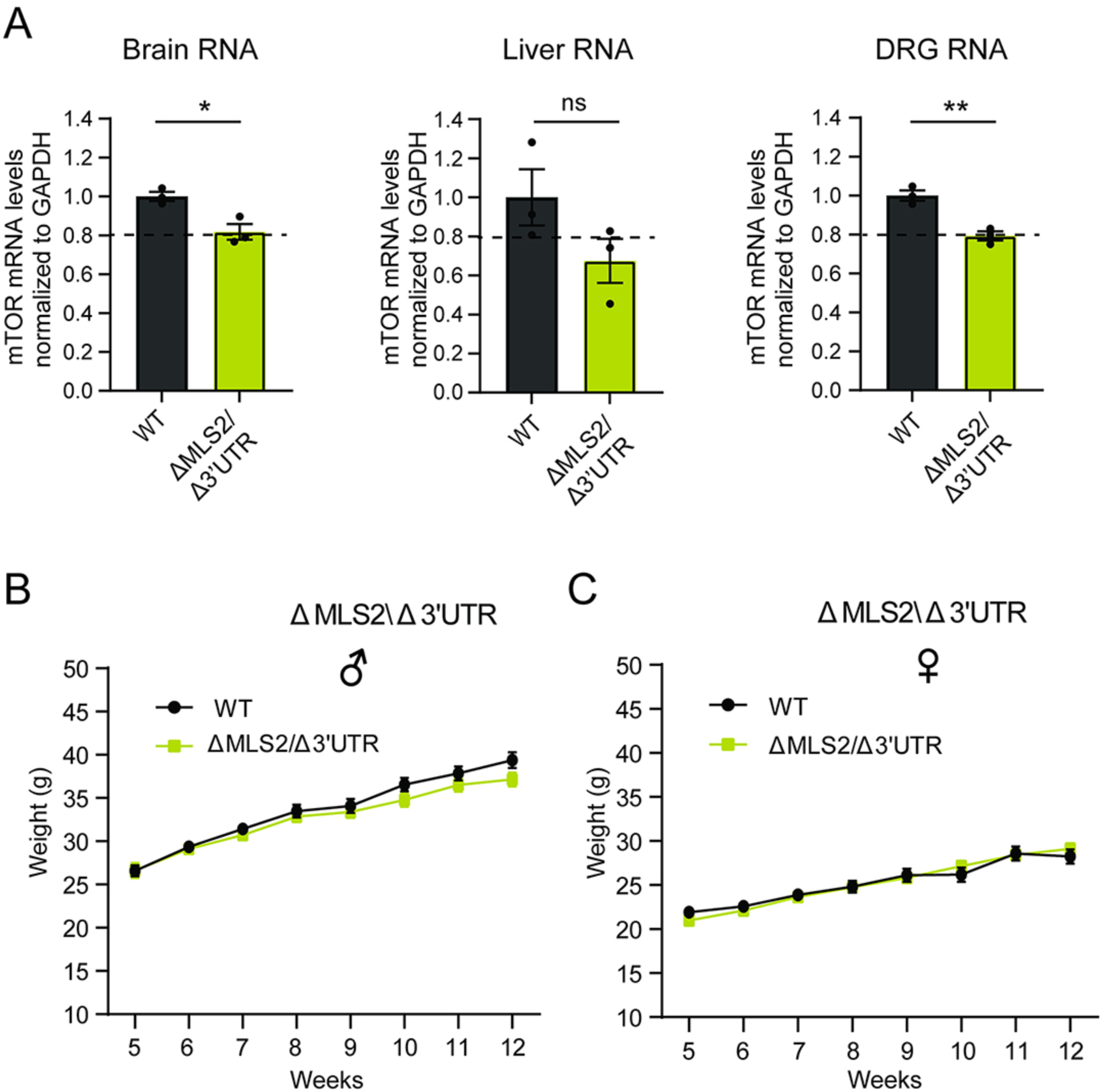
Further characterization of the mTOR ΔMLS2/Δ3’UTR gene-edited mouse line. (**A**) mTOR mRNA levels in brain, liver and DRG, analyzed by RT-qPCR, n≥3; *p<0.05, **p<0.01, two tails unpaired t-test. (**B**, **C**) Growth and size of mTOR ΔMLS2/Δ3’UTR male (**B**) and female (**C**) mice assessed by weekly weight measurements over the indicated time frame. Data represent mean ± SEM, n ≥ 11, Two-way ANOVA with Tukey’s post-hoc test shows no significant differences between genotypes.

## Notes

### Competing Interest Statement

The authors have declared no competing interest.

